# The small molecule CBR-5884 inhibits the *Candida albicans* phosphatidylserine synthase

**DOI:** 10.1101/2023.11.01.565123

**Authors:** Yue Zhou, Gregory A. Phelps, Mikayla M. Mangrum, Jemma McLeish, Elise K. Phillips, Jinchao Lou, Christelle F. Ancajas, Jeffrey M. Rybak, Peter M. Oelkers, Richard E. Lee, Michael D. Best, Todd B. Reynolds

## Abstract

Systemic infections by *Candida spp.* are associated with high mortality rates, partly due to limitations in current antifungals, highlighting the need for novel drugs and drug targets. The fungal phosphatidylserine synthase, Cho1, from *Candida albicans* is a logical antifungal drug target due to its importance in virulence, absence in the host and conservation among fungal pathogens. Inhibitors of Cho1 could serve as lead compounds for drug development, so we developed a target-based screen for inhibitors of purified Cho1. This enzyme condenses serine and cytidyldiphosphate-diacylglycerol (CDP-DAG) into phosphatidylserine (PS) and releases cytidylmonophosphate (CMP). Accordingly, we developed an *in vitro* nucleotidase-coupled malachite green-based high throughput assay for purified *C. albicans* Cho1 that monitors CMP production as a proxy for PS synthesis. Over 7,300 molecules curated from repurposing chemical libraries were interrogated in primary and dose-responsivity assays using this platform. The screen had a promising average Z’ score of ∼0.8, and seven compounds were identified that inhibit Cho1. Three of these, ebselen, LOC14, and CBR-5884, exhibited inhibitory effects against *C. albicans* cells, with fungicidal inhibition by ebselen and fungistatic inhibition by LOC14 and CBR-5884. Only CBR-5884 showed evidence of disrupting *in vivo* Cho1 function by inducing phenotypes consistent with the *cho1*ΔΔ mutant, including a reduction of cellular PS levels. Kinetics curves and computational docking indicate that CBR-5884 competes with serine for binding to Cho1 with a *K*_i_ of 1,550 ± 245.6 nM. Thus, this compound has the potential for development into an antifungal compound.

## Introduction

*Candida* species are the most commonly isolated fungal pathogens of humans (1, 2). *Candida* has been associated with mucosal infections, such as vulvovaginal infections in 51% of women or oropharyngeal infections in 27% of HIV+ patients, even when on anti-retroviral (ART) therapy (3, 4). In addition, these species are responsible for ∼27% of bloodstream infections associated with a central line (1, 2, 5). *Candida* bloodstream infections pose a considerable threat to public health, with a mortality rate of approximately 40% (1, 2, 6, 7). *Candida albicans* is the most commonly isolated species within the *Candida* genus (1, 2, 8). Currently, there are only three classes of antifungal drugs in common use (azoles, polyenes, and echinocandins) for treating systemic *Candida* infections. However, their effectiveness is hindered by rising drug resistance to azoles and echinocandins and the toxicity profile of the polyene amphotericin B (9–13).

Therefore, there is a pressing need to develop novel antifungal drugs.

Central in the phospholipid synthetic pathway in fungi is the phosphatidylserine (PS) synthase reaction (CDP-diacylglycerol—serine O*-*phosphatidyltransferase; EC 2.7.8.8) mediated by Cho1. This enzyme has been identified as a potential drug target due to the observation that [i] disruption of Cho1 in *C. albicans* prevents this fungus from causing disease in mouse models of systemic or oral infection (14, 15), and this enzyme is also crucial for the growth of the major fungal pathogen *Cryptococcus neoformans* (16); [ii] Cho1 is not present in humans, indicating that specific inhibitors targeting this enzyme should not have toxic effects on humans (17), and [iii] *CHO1* is highly conserved across various fungal species, suggesting that inhibitors of this enzyme would have broad spectrum anti-fungal effects (16, 17). Hence, inhibitors of Cho1 would be excellent lead compounds for antifungal drug development.

The fungal Cho1 enzyme was first characterized in the yeast *Saccharomyces cerevisiae*. This included descriptions of cellular localization in the endoplasmic reticulum and mitochondrial outer membranes (18–20), activity regulation (21–23) and protein purification (24, 25). Cho1 catalyzes the formation of PS from cytidyldiphosphate-diacylglycerol (CDP-DAG) and L-serine, and it belongs to the <u>CDP</u>- <u>a</u>lcohol <u>p</u>hosphatidyltransferase (CDP-AP) protein family. The CDP-AP proteins employ the highly conserved <u>C</u>DP-<u>a</u>lcohol <u>p</u>hosphotransferase (CAPT) motif, D-(X)_2_-D-G-(X)_2_- A-R-(X)_2_-N-(X)_5_-G-(X)_2_-L-D-(X)_3_-D, to bind CDP-linked molecules and facilitate the formation of a phosphodiester bond between the CDP-linked molecule and another small alcohol (16, 26–30), specifically, CDP-DAG and serine for Cho1. However, it is important to note that the binding pocket for serine, unlike the CAPT motif, is not conserved among CDP-AP proteins. Previous studies have identified and characterized several crucial residues within the CAPT motif and the putative serine-binding site of *C. albicans* Cho1 through alanine scanning mutagenesis (26, 31). Additionally, valuable insights have been provided into the serine-binding pocket from the atomic structure of the PS synthase from the archaean *Methanocaldococcus jannaschii* (32). Differences between the *M. jannaschii* PS synthase and *C. albicans* Cho1 are apparent by the presence of specific residues in *C. albicans* Cho1 that play important roles, yet lack clear counterparts in the *M. jannaschii* PS synthase (26, 32).

Besides the characterization of the substrate-binding residues, the *C. albicans* Cho1 has also been solubilized and purified as a hexameric protein (33), distinct from all the CDP-AP enzymes with solved structures (30, 32, 34–40), which are dimers. The hexameric *C. albicans* Cho1 can be separated into a trimer of stable dimers, indicating the hexamer might be [i] an early oligomer state, since Cho1 was solubilized from the early-to-mid log phase of *C. albicans* or [ii] species-specific (33). Furthermore, purified Cho1 enzyme was optimized for activity and was shown to have a *K*_m_ for CDP-DAG of 72.20 μM with a *V*_max_ of 0.079 nmol/(μg*min) while exhibiting a sigmoidal kinetic curve for its other substrate serine, indicating cooperative binding (33). This sigmoidal kinetic could potentially reconcile the contradicting high and low *K*_m_ values reported previously for *S. cerevisiae* PS synthase (22, 25, 41–43). The mechanism underlying the cooperative binding of serine is currently unknown.

Rational drug design is one way to discover compounds to a drug target protein. This can be achieved through either ligand-based or structure-based design methods (44). However, since there is a scarcity of known Cho1 ligands, and the atomic structure of *C. albicans* Cho1 has not yet been solved, employing rational drug design to identify Cho1- specific inhibitors is challenging. On the contrary, small molecule screening is an alternative way to identify inhibitors to Cho1 independent of structural information. Two whole-cell screens have been carried out to identify Cho1 inhibitors, but neither has been successful (45, 46). One identified the compound SB-224289, but it was discovered that SB-224289 only affects Cho1-associated physiological pathways (45). The other screen identified bleomycin, but this again impacts phospholipid related physiologies rather than Cho1 itself (46). Thus, to carry out identification of a Cho1 inhibitor, a target-based screen was developed. This approach is enabled by the purification of Cho1, and is favorable because molecules identified will show direct inhibition on the target. Potential issues such as the cellular entry of molecules identified from target-based screening can be resolved later through medicinal chemistry approaches.

Cho1 activity has been measured in crude membrane preps (26, 31, 45, 47) and as a purified form (33, 48), using a radioactive substrate. However, that methodology is inconsistent with a high throughput screen. Here, we have adapted a non-radioactive assay with an easy setup and colorimetric readout (49), which detects the byproduct cytidine monophosphate (CMP) released from Cho1, to measure its activity in the presence of screening molecules. Using this assay, approximately 7,300 molecules were interrogated in a primary screen from a set of curated repurposing libraries to reveal one compound, CBR-5884, that stood out as it displayed an inhibitory effect on Cho1 both *in vitro* and in live *C. albicans* cells.

## Results

### A malachite-green-based nucleotidase-coupled assay was used to screen for inhibitors of purified Cho1 protein

A expression cassette plasmid carrying the strong, constitutive promoter for translational elongation factor 1 (*P_TEF1_*) fused upstream of the *C. albicans CHO1* gene was integrated into the genome at the *TEF1* locus to ensure a strong and stable expression level (50). Then, Cho1 was solubilized and purified from the microsomal fraction of *Candida albicans* as described in the Materials and Methods section. A blue native PAGE indicated that the Cho1 protein was purified to relative homogeneity as a hexameric form of ∼180 kDa (Figure 1A), consistent with findings in (33). The purified Cho1 was used for small molecule screening.

**Figure 1.**
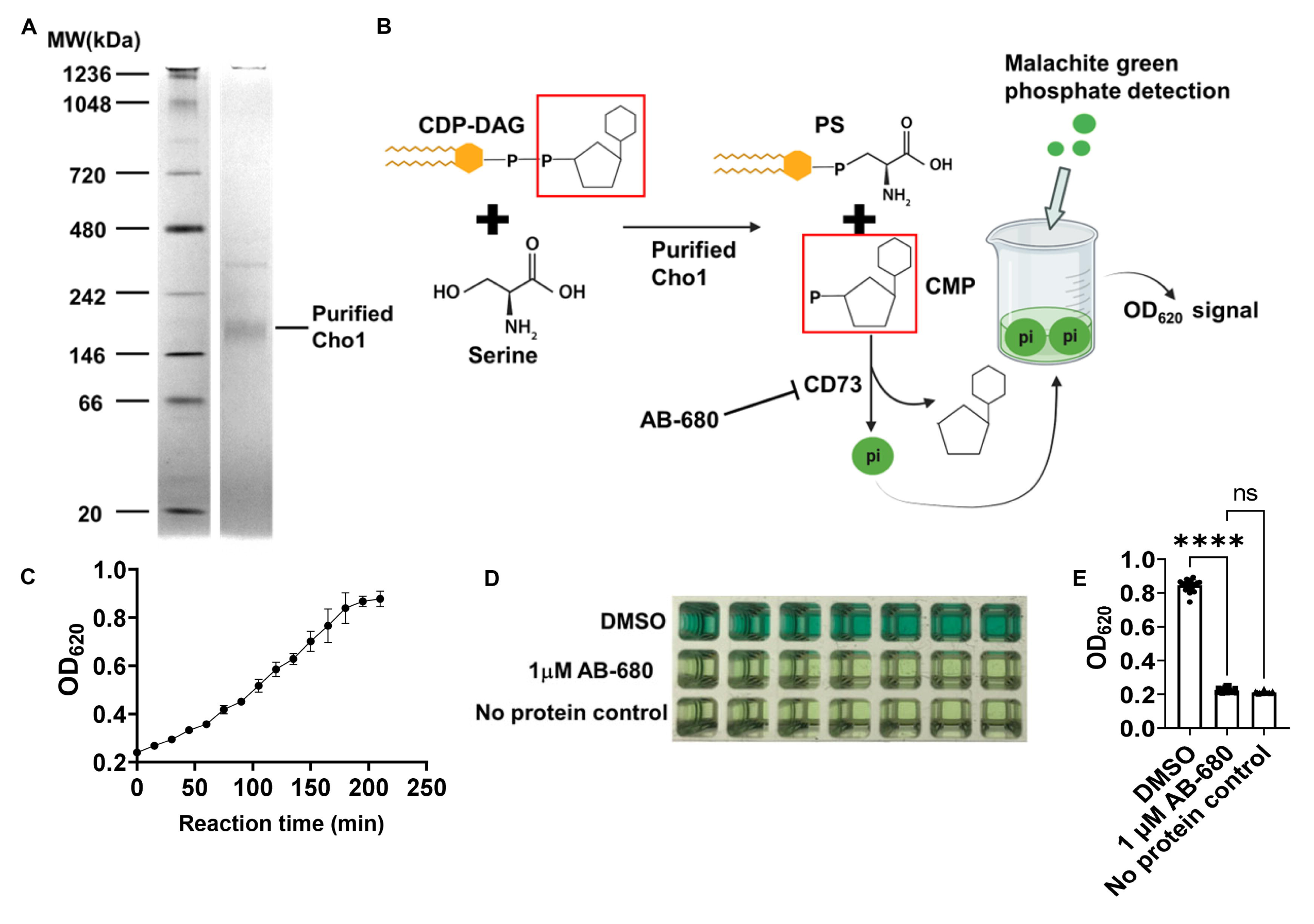
A malachite-green-based nucleotidase-coupled assay measures the activity of purified Cho1 protein. (A) Blue native PAGE gel of the purified hexameric tag-free Cho1 protein. Purified Cho1 and protein ladder with known MW are indicated. The gel was stained with Coomassie Blue R-250. **(B)** Schematic representation of the malachite-green-based nucleotidase-couple assay. Cho1 synthesizes PS from CDP-DAG (cytidyldiphosphate-diacylglycerol) and serine. This releases PS and CMP (cytidylmonophosphate). The phosphate from CMP is cleaved by the nucleotidase CD73 to release inorganic phosphate, which can be bound by the malachite green reagent and measured colorimetrically at OD_620_. AB-680 is a potent inhibitor of CD73, and can thus inhibit the reaction. **(C)** OD_620_ signal from the malachite green reagent that was added to the reaction shown in **(B)** at different time points after the reaction started. Reactions were set up with the same conditions and stopped by adding malachite green at the time indicated. The dots represent the mean of four replicates, and the error bars are ± standard deviation (S.D.) values. **(D)** Inhibition of the nucleotidase-coupled assay by AB-680 is shown for a series of replicates and **(E)** is quantified for a total of 21 replicates. Statistics were conducted using one-way ANOVA using Tukey’s multiple comparisons test (ns=not significant, ****, p < 0.0001).

A malachite green-based nucleotidase-coupled assay was used to measure the PS synthesis activity of Cho1 (Figure 1B). Cho1 catalyzes the production of PS and cytidylmonophosphate (CMP) from CDP-DAG and serine, where CMP can then be recognized and cleaved by the nucleotidase CD73 to release inorganic phosphate, which is subsequently detected by malachite green. To test whether this malachite green-based assay can reflect Cho1 activity, a time-course test was performed with a fixed amount of purified Cho1 in the presence of substrates (Figure 1C). The OD_620_ signal increased as the reaction proceeded until it plateaued at 200 min, indicating that the assay is suitably dynamic to probe Cho1 activity. Based on this data, a fixed time of 180 minutes was chosen for the screen.

As no effective inhibitors of Cho1 have been identified to date, a highly potent and selective inhibitor of the second step of the assay (nucleotidase CD73) was examined as a positive control (51). This compound, AB-680, exhibited an IC_50_ value of 1.82 nM in the malachite-green assay (Figure S1A). A concentration of 1 μM AB-680 was used in the screen, and was compared to DMSO and no protein wells, which served as negative controls. Reactions with AB-680 showed similar color as the no protein control in the 384-well plate (Figure 1D), and the measured OD_620_ signal was four-fold less in the presence of the AB-680 (Figure 1E). Thus, AB-680 can be used as a positive control for identifying inhibitors in the primary screen. In order to eliminate false positives that are actually inhibiting the nucleotidase CD73, a counter screen was developed where Cho1 was replaced with CMP, but CD73 was still present so that we could detect direct inhibitors of the nucleotidase.

### Seven Cho1-specific inhibitors were identified from the high throughput small molecule screen

The screen interrogated 7,307 molecules from three curated repurposing libraries in the primary screen and counter screen (which were run concurrently) at a final concentration of 100 μM each (Figure 2A). The primary screen had an average Z’ score of ∼0.8 (Figure S1B), indicating a good signal to noise ratio (52). To prioritize hit compounds and eliminate false positives, % Δinhibition (Figure 2B) was calculated by subtracting the % inhibition from the counter screen (Figure S1D) from that of the primary screen (Figure S1C) for each molecule. All compounds from the >80% Δinhibition population and selected molecules with limited structural liabilities from the 50-80% Δinhibition population (for a total of 82 molecules) were advanced for dose-response assessment using the same screening platform, which once again yielded results of high quality (Z’ of 0.83; Figure S1E). Compounds exerting dose-dependent activity were then further triaged by manual inspection to exclude <u>p</u>an <u>a</u>ssay <u>in</u>terference compound<u>s</u> (PAINS), which tend to react nonspecifically with numerous biological targets (53). Finally, seven molecules exerting IC_50_ values ≤ 76 μM were identified, of which CBR-5884, ML-345, ebselen and tideglusib were the most potent, possessing IC_50_ ≤ 20 μM (Figure 2C). These molecules, in addition to avasimibe and LOC14 were selected for further investigation. The TC-N 22A molecule was not easily available and was not pursued.

**Figure 2.**
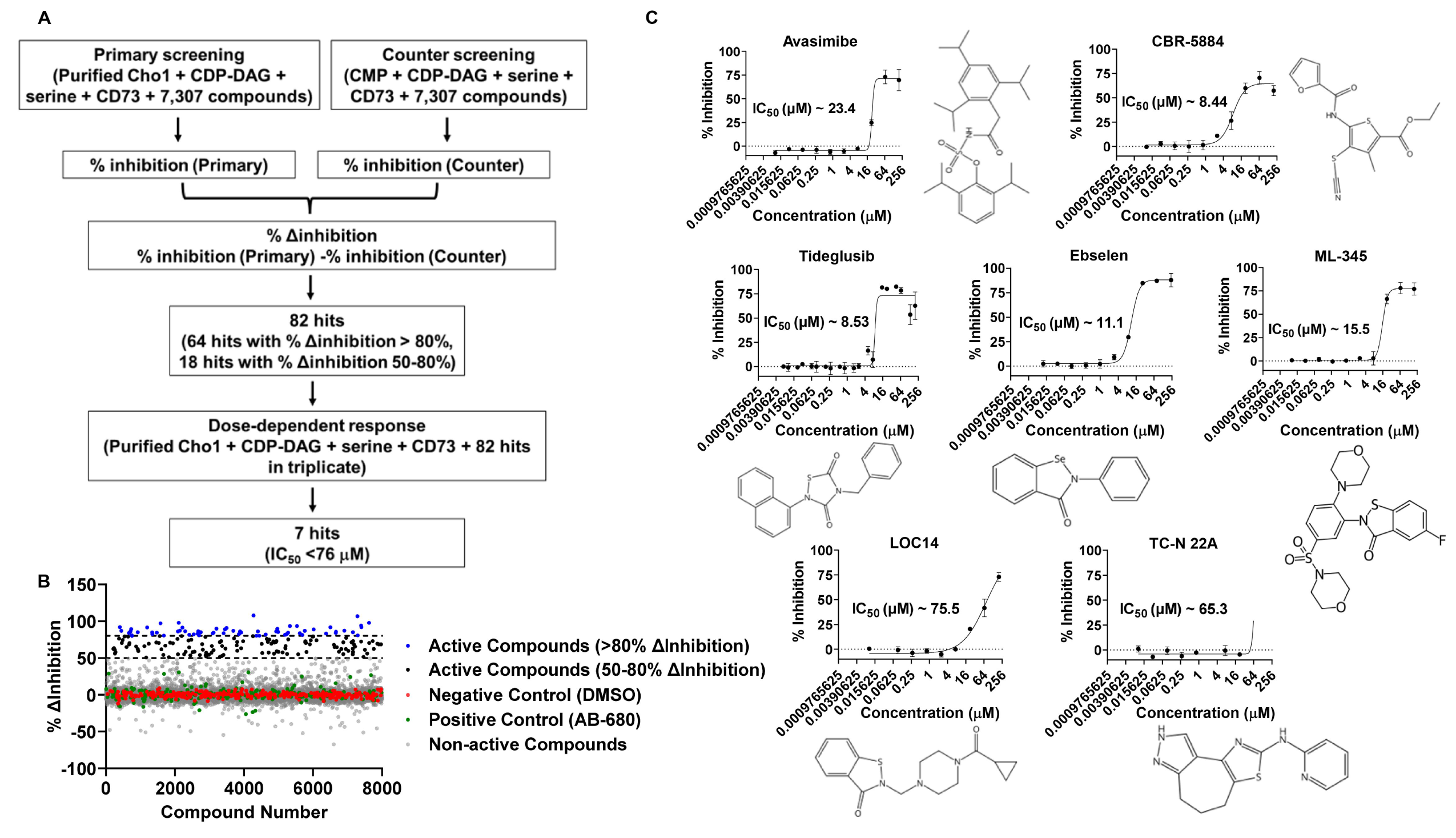
Seven Cho1-specific inhibitors were identified from the high-throughput malachite green screen. (A) Flowchart for the primary and counter screen and the calculation of % Δinhibition. **(B)** The dot plot of % Δinhibition for all the compounds, including controls, used in the screen. Reaction with DMSO and AB-680 were used as 0% Δinhibition (negative) and 100% Δinhibition (positive) controls, respectively. The two dotted lines from the Y-axis indicate 80 and 50% Δinhibition, respectively. **(C)** Dose-response curve and structure of the seven non-pan-assay interference (non-PAINS) compounds identified from the screen. The dots represent the mean of three replicates, and the error bars are ± standard deviation (S.D.) values. Best-fit IC_50_ values (in μM) were shown in each graph.

### Ebselen, LOC14 and CBR-5884 showed inhibitory effects on *C. albicans* cells

To further validate the inhibitory effects of the six molecules on Cho1, a radioactive PS synthase assay was conducted on Cho1 in the presence of these compounds. Unlike the malachite green-based assay, the radioactive PS synthase assay directly measures the incorporation of L-[^3^H]-serine into PS in the lipid phase (26, 33, 47). The radioactive PS synthase assay was performed on purified Cho1 in the presence of the six compounds at 100 μM, and all six molecules were shown to totally inhibit Cho1 (Figure 3A), consistent with the screening results.

**Figure 3.**
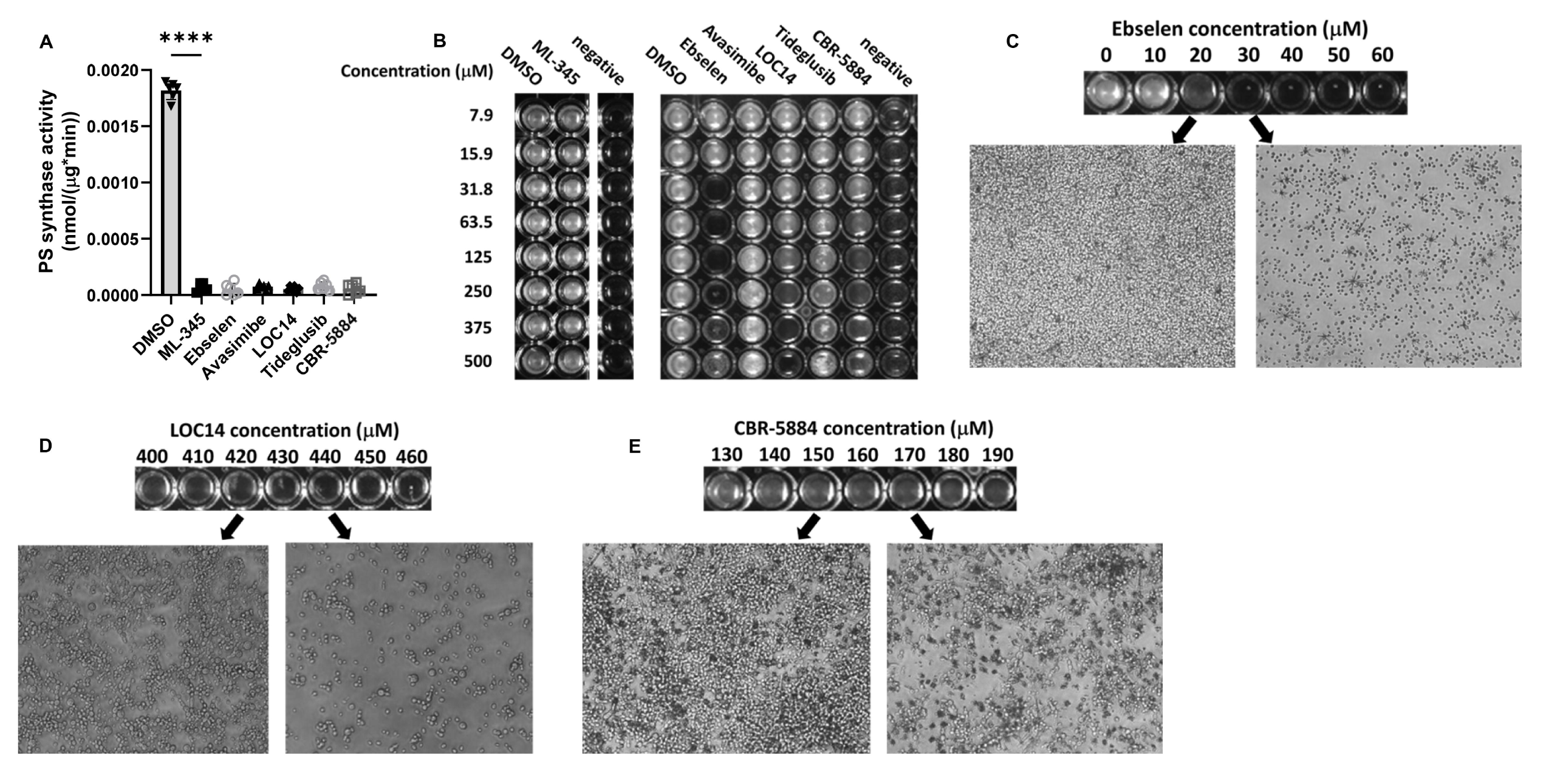
Ebselen, LOC14 and CBR-5884 inhibited cell growth of *C. albicans*. (A) The PS synthase activity of purified Cho1 was measured by L-[^3^H]-serine incorporation into PS in the presence of different inhibitors at 100 μM or equivalent DMSO, and are presented as nmol/(μg protein*min). Statistics were conducted using one-way ANOVA and Dunnett’s T3 multiple comparisons test (****, 0.0001> p). The activities were measured in duplicate with a total of six biological replicates as indicated. The bars represent the mean and the error bars are ± S.D. values. **(B)** Wildtype *C. albicans* was grown in minimal medium for 24 hours with DMSO or different inhibitors at the concentrations indicated, and the *cho1*ΔΔ strain in DMSO was used as a control (shown as “negative”). **(C-E)** Inverted microscopy (400x) was used to visualize cells treated with **(C)** ebselen **(D)** LOC14 and **(E)** CBR-5884 under the indicated concentrations for 24 hours. Black arrows indicate from which concentrations of compounds the images are taken.

We then tested the effects of the six compounds on live cells. PS is the lipid precursor for making phosphatidylethanolamine (PE), an essential phospholipid, via the *de novo* pathway. Growth can resume when PS synthesis is inhibited, albeit more slowly, if the organism is able to make PE from ethanolamine acquired from the medium via the salvage (Kennedy) pathway (14, 26). Thus, ethanolamine-dependent growth in minimal media is a characteristic phenotype of PS synthesis loss.

We first characterized the inhibitory effects of the compounds on live cells in minimal media lacking ethanolamine. Wildtype *C. albicans* strain SC5314 was grown with avasimibe, CBR-5884, tideglusib, ebselen, ML-345 and LOC14 in minimal media at 30°C for 24 hours, along with a *cho1*ΔΔ strain as a control (Figure 3B). The concentration of the compounds was varied from 7.9 to 500 μM. Avasimibe, tideglusib and CBR-5884 precipitated at high concentrations, indicating a low solubility (Figure S2A). Among all compounds, avasimibe, tideglusib and ML-345 did not affect cell growth, even at 500 μM, indicating they do not have an inhibitory effect on *C. albicans* cells under our assay conditions (Figure 3B). Ebselen, LOC14 and CBR-5884, on the contrary, caused cell growth perturbation at different concentrations (Figure 3B). This is consistent with the radioactive PS synthase assay done on the crude membrane containing Cho1, in which only ebselen, CBR-5884 and LOC14 decreased PS production on the native crude membranes (Figure S3).

To more precisely define the inhibitory concentrations of ebselen, LOC14 and CBR-5884 on live cells, wildtype *C. abicans* SC5314 was incubated with these compounds in a dosage series with 10 μM increments to determine a more accurate minimal inhibitory concentration (MIC) value. Ebselen at 30 μM was shown to inhibit cell growth, and cells were smaller than normal wild-type cells when viewed in the microscope (Figure 3C). LOC14 and CBR-5884, on the contrary, caused significantly decreased cell densities at 440 μM and 170 μM, respectively (Figures 3D&3E), but not abnormally small cells. To determine if the inhibitory effects were fungistatic or fungicidal, cells grown for 24 hours in minimal media with these compounds were plated, and colony forming units (CFUs) were counted. Ebselen showed no colonies after incubation at 30 μM, which is consistent with a fungicidal effect (Figure 4A). In contrast, cells incubated with 430 μM LOC14 and 170 μM CBR-5884 exhibited similar CFUs as the pre-incubation, indicative of a fungistatic effect (Figures 4B&C).

**Figure 4.**
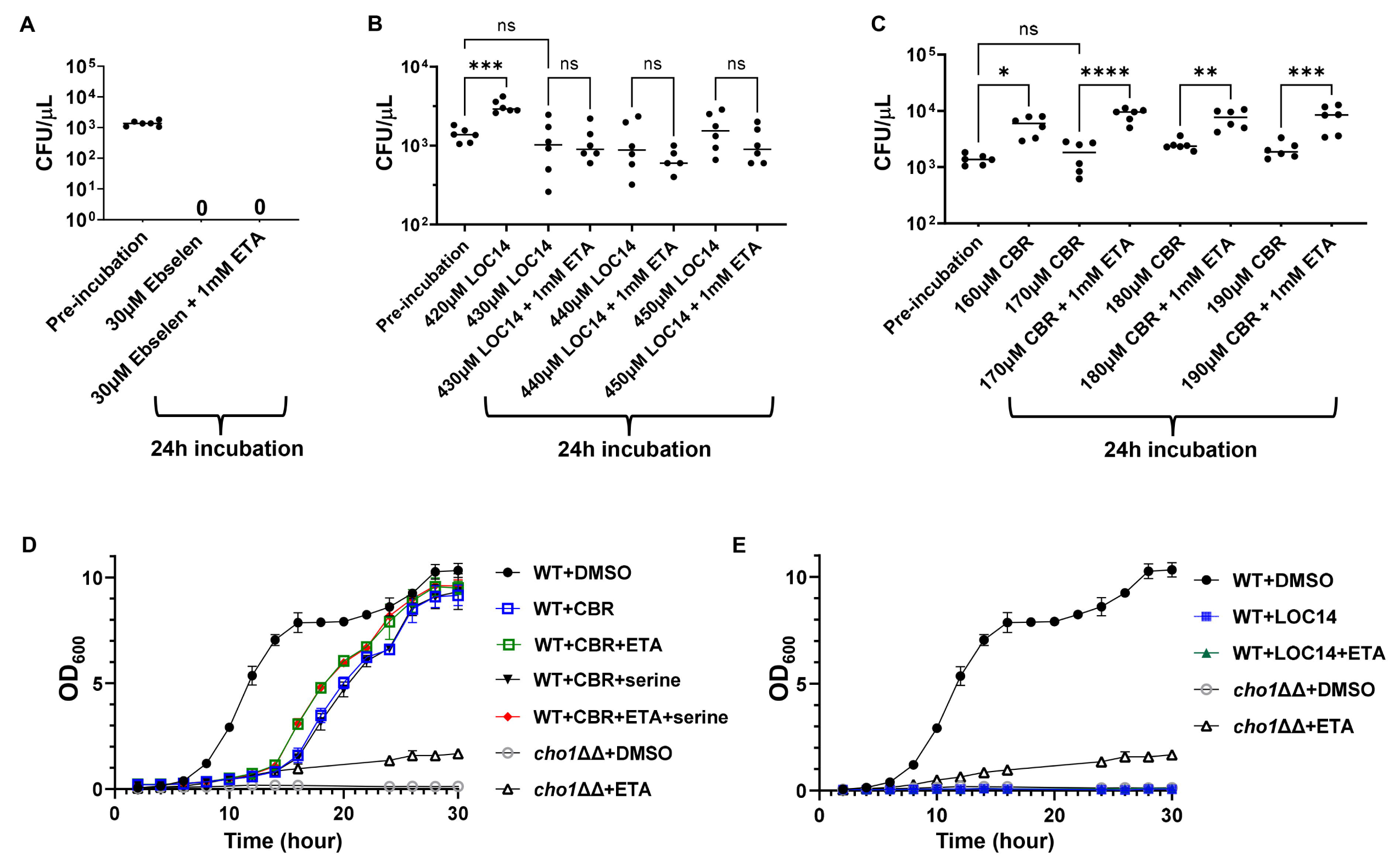
Ethanolamine supplementation can mitigate the inhibitory effects of CBR- 5884. (A-C) Colony forming units (CFUs) from cells treated with different concentrations of **(A)** ebselen, **(B)** LOC14 and **(C)** CBR-5884, ± 1 mM ethanolamine (ETA), for 24 hours compared to pre-incubation. Statistics were conducted using one-way ANOVA and Tukey’s multiple comparisons test (ns=not significant, p > 0.05; *, 0.05> p > 0.01, **, 0.01> p > 0.001). Six biological replicates were tested in each treatment. **(D)** Growth curves of wildtype *C. albicans* in the presence of 170 μM CBR- 5884 or equivalent DMSO, with the addition of 1 mM ethanolamine (ETA) or 5 mM serine or both, from 0 to 30 hours. The *cho1*ΔΔ strain was also included as a control. The dots represent the mean values of six replicates, and the error bars are ± standard deviation (S.D.) values. **(E)** Growth curves of wildtype *C. albicans* in the presence of 430 μM LOC-5884 or equivalent DMSO, with the addition of 1 mM ethanolamine (ETA), from 0 to 30 hours. The *cho1*ΔΔ strain was also included as a control. The dots represent the mean values of six replicates, and the error bars are ± standard deviation (S.D.) values.

### The inhibitory effect of CBR-5884 can be rescued by ethanolamine supplementation

To determine if the impacts of ebselen, LOC14 and CBR-5884 on cells were consistent with perturbed Cho1 function, we repeated the growth inhibition assay at the optimized drug concentrations plus or minus supplementation with 1mM ethanolamine. Although ethanolamine did not rescue cells with ebselen or LOC14, cells treated with the MIC (≥170µM) of CBR-5884 generated significantly higher CFUs upon addition of ethanolamine (Figures 4A-C). This result suggests that the inhibitory effect of ebselen and LOC14 on the cells are not solely caused by Cho1 inhibition, while CBR-5884 is more directly targeting Cho1.

To gain insight into the dynamic, inhibitory properties of these molecules, growth curves were determined with wildtype cells grown in the minimal media ± ethanolamine, in the presence of CBR-5884 or LOC14. Ebselen was not pursued in the growth curve assay due to its fungicidal effect. The *cho1*ΔΔ strain was used as a control for loss of Cho1 activity. Given that CBR-5884 is a selective inhibitor of phosphoglycerate dehydrogenase that is involved in *de novo* serine synthesis in cancer cells (54), and to make sure that the growth perturbation is not due to serine starvation, 5 mM serine was also added to the media ± ethanolamine. As shown in Figure 4D, cells grown in media + ethanolamine and + ethanolamine/serine grew similarly, while those grown in minimal media and minimal media + serine had similarly reduced growth. This suggests that serine did not help the cells recover from the inhibition from CBR-5884, which likely indicates [i] CBR-5884 does not target *C.albicans* phosphoglycerate dehydrogenase or [ii] the CBR-5884 inhibition of *C. albicans* phosphoglycerate dehydrogenase is not the major cause of diminished growth. Cells with ethanolamine supplementation grew better than those in minimal media alone, especially from 12 to 24 hours, consistent with Figure 4C. Growth rates during log phase (Figure S2B) and lag phase duration (Figure S2C) were estimated from the growth curves, and they showed that the addition of ethanolamine increased the growth rate and decreased the lag time. All of these results again support the hypothesis that CBR-5884 targets Cho1 *in vivo*. All strain groups caught up with growth after a 24-hour incubation (Figure 4D), suggesting that CBR-5884 loses its effect over time.

In contrast, LOC14 was also subjected to growth curve determination but no cells grew, in the presence or absence of ethanolamine (Figure 4E). This corroborates with Figure 4B that the inhibition by LOC14 is not acting solely on Cho1. Since the goal was a Cho1-specific inhibitor, ebselen and LOC14 were not pursued further.

### CBR-5884 interferes with PS synthesis *in vivo*

It has been determined previously that deletion of Cho1 leads to increased exposure (unmasking) of cell wall β(1–3)-glucan, rendering cells are more prone to be targeted by the immune system (14, 55). Here, we tested whether CBR-5884 could induce unmasking. Wildtype *C. albicans* cells were grown in rich medium (YPD) supplemented with DMSO or CBR-5884 for 30, 60 and 120 min, along with the *cho1*ΔΔ strain control, and exposed β(1–3)-glucan was stained and visualized through confocal microscopy. The *cho1*ΔΔ strain exhibited increased unmasked foci, compared to the WT strain in DMSO, consistent with previous findings that disruption of Cho1 leads to increased unmasking (Figure 5A) (55). Similarly, CBR-5884 treatment showed increased unmasking after 30-min incubations, and the unmasking became more obvious at 60- and 120-min treatment. The mean fluorescence values confirmed that 30-min CBR-5884 treatment is sufficient to induce significantly increased unmasking compared to wildtype *C. albicans*, and 120-min treatment could induce more unmasking than the *cho1*ΔΔ mutant (Figure 5B). This could be because CBR-5884 inhibition causes a sudden loss of Cho1 function that the cells have not yet adjusted to, whereas a *cho1*ΔΔ mutant has adjusted its metabolism to the loss of PS. Alternatively, it may indicate an off-target effect. It is also interesting to note that there are dark structures in the cells under bright field microscopy when treated with CBR-5884, which are absent in wildtype or *cho1*ΔΔ strains without CBR-5884. We currently cannot explain the identity or formation of these structures, but speculate that they represent fragmented vacuoles that serve in detoxification (56).

**Figure 5.**
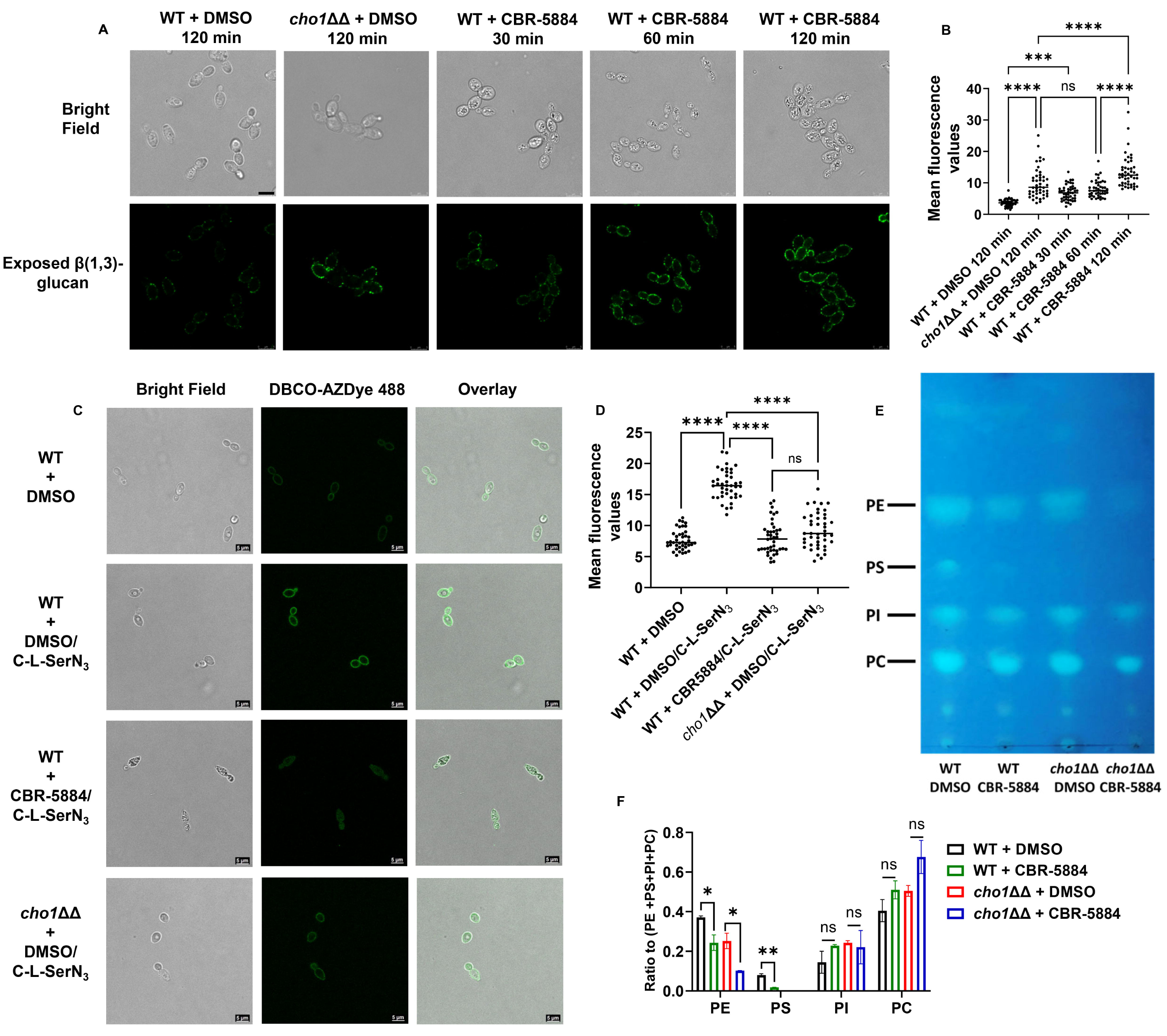
CBR-5884 interferes with *in vivo* PS synthesis. (A) CBR-5884 induces cell wall β(1,3)-glucan exposure. Exposed cell wall β(1,3)-glucan of wildtype *C. albicans* treated with 170 μM CBR-5884 or equivalent DMSO control for times indicated are shown. Exposed β(1,3)-glucan is shown as green fluorescence and the corresponding cells are shown in the bright field as well. **(B)** Quantification of the mean fluorescence from **(A)**. Forty-six cells from at least 10 images were used for the quantification. Statistics were conducted using one-way ANOVA and Tukey’s multiple comparisons test (ns=not significant, p > 0.05; ****, 0.0001> p). **(C)** CBR-5884 interferes with the incorporation of C-L-SerN_3_ probe into the cell membrane. A final concentration of 1.5 mM C-L-SerN_3_ was added to wildtype *C. albicans* and *cho1*ΔΔ cells grown with and without 170 μM CBR-5884. The cells were then stained with the DBCO-AZ Dye 488 to allow click-tagging, followed by microscopy. Corresponding brightfield and overlay images were also shown. Wildtype *C. albicans* grown without the C-L-SerN_3_ probe was included as a control for background fluorescence. **(D)** Quantification of the mean fluorescence from **(C)**. Forty cells from at least 10 images were used for the quantification. Statistics were conducted using one-way ANOVA and Tukey’s multiple comparisons test (ns=not significant, p > 0.05; ****, 0.0001> p). **(E)** Thin layer chromatography (TLC) plate of phospholipids extracted from wildtype and *cho1*ΔΔ *C. albicans* treated with 170 μM CBR-5884 or equivalent DMSO. The positions of PS (phosphatidylserine), PE (phosphatidylethanolamine), PI (phosphatidylinositol) and PC (phosphatidylcholine) are indicated based on standards. **(F)** The ratio of the phospholipid to total (PE+PS+PI+PC) phospholipids for strains in **(E)**. The quantification was done in ImageJ software from two TLC plates. Statistics were conducted using unpaired two-tailed t test (ns=not significant, p > 0.05; *, 0.05> p > 0.01, **, 0.01> p > 0.001).

In order to more directly measure the impact of CBR-5884 on PS synthesis *in vivo*, an assay for fluorescence-based labeling of PS via biorthogonal tagging using a clickable serine probe was utilized. This probe consists of a serine analogue carrying an azide tag for click chemistry, C-L-SerN_3_. This probe has been demonstrated to be incorporated into live cell membranes by infiltrating lipid metabolism to produce azide-tagged PS analogues that can be post-labeled by click-tagging with fluorophores to localized/quantify PS in cells (57, 58). Wildtype *C. albicans* was grown in YPD to early log phase before C-L-SerN_3_ was added, along with CBR-5884 or DMSO (Figure 5C).

The *cho1*ΔΔ strain and no probe controls were also included for background fluorescence, and mean fluorescence of each group was quantified (Figure 5D). In the absence of CBR- 5884, C-L-SerN_3_ labels the cell membrane of wildtype *C. albicans* with stronger fluorescence compared to no probe or *cho1*ΔΔ strains (Figures 5C&5D), indicating C-L- SerN3 was converted into PS as previously described (57). However, in the presence of CBR-5884, the fluorescence is significantly diminished on the periphery of the cell (Figures 5C&D). This indicates that CBR-5884 inhibits *in vivo* PS production.

Finally, a direct biochemical test for PS levels was performed by thin layer chromatography (TLC) in cells treated with CBR-5884. Wildtype *C. albicans* and *cho1*ΔΔ strains were grown in the presence and absence of CBR-5884, and the four major phospholipid species are shown in Figure 5E and quantified in Figure 5F. Consistently, the relative PS level in wildtype *C. albicans* strain treated with CBR-5884 significantly dropped compared to the DMSO treatment (Figures 5E&5F), indicating that CBR-5884 interferes with PS production. Interestingly, the relative PE levels also significantly decreased in strains treated with CBR-5884, especially in the *cho1*ΔΔ strain without PS as a precursor (Figures 5E&5F). This potentially indicates that CBR-5884 may independently impact PE production.

### CBR-5884 acts as a competitive inhibitor that occupies the serine binding site of Cho1

To investigate the molecular mechanism by which CBR-5884 inhibits Cho1, purified Cho1 specific activity was assayed with varying concentrations of serine and CDP-DAG in the presence of CBR-5884. Serine was varied from 4 to 32 mM, CDP-DAG was kept at 200 μM, and several concentrations of CBR-5884 were tested (Figure 6A). The pattern of inhibition fits with competitive inhibition and a low *K*_i_ value of 1,550 ± 245.6 nM. Next, serine was held at a sub-saturating concentration of 20 mM or a saturating concentration of 32 mM, and CDP-DAG was varied from 25 to 300µM. At the lower serine concentration, the inhibition of CBR-5884 on Cho1 activity was observed (Figure 6B), but the inhibition was overcome under saturating serine concentrations (Figure 6C). These results suggest that CBR-5884 inhibits Cho1 by competing for serine and can be outcompeted with a high serine concentration. It is interesting that CDP-DAG inhibits Cho1 activity at high concentrations, especially in the presence of CBR-5884 (Figure 6B). This substrate inhibition from CDP-DAG has been reported previously (22, 59).

**Figure 6.**
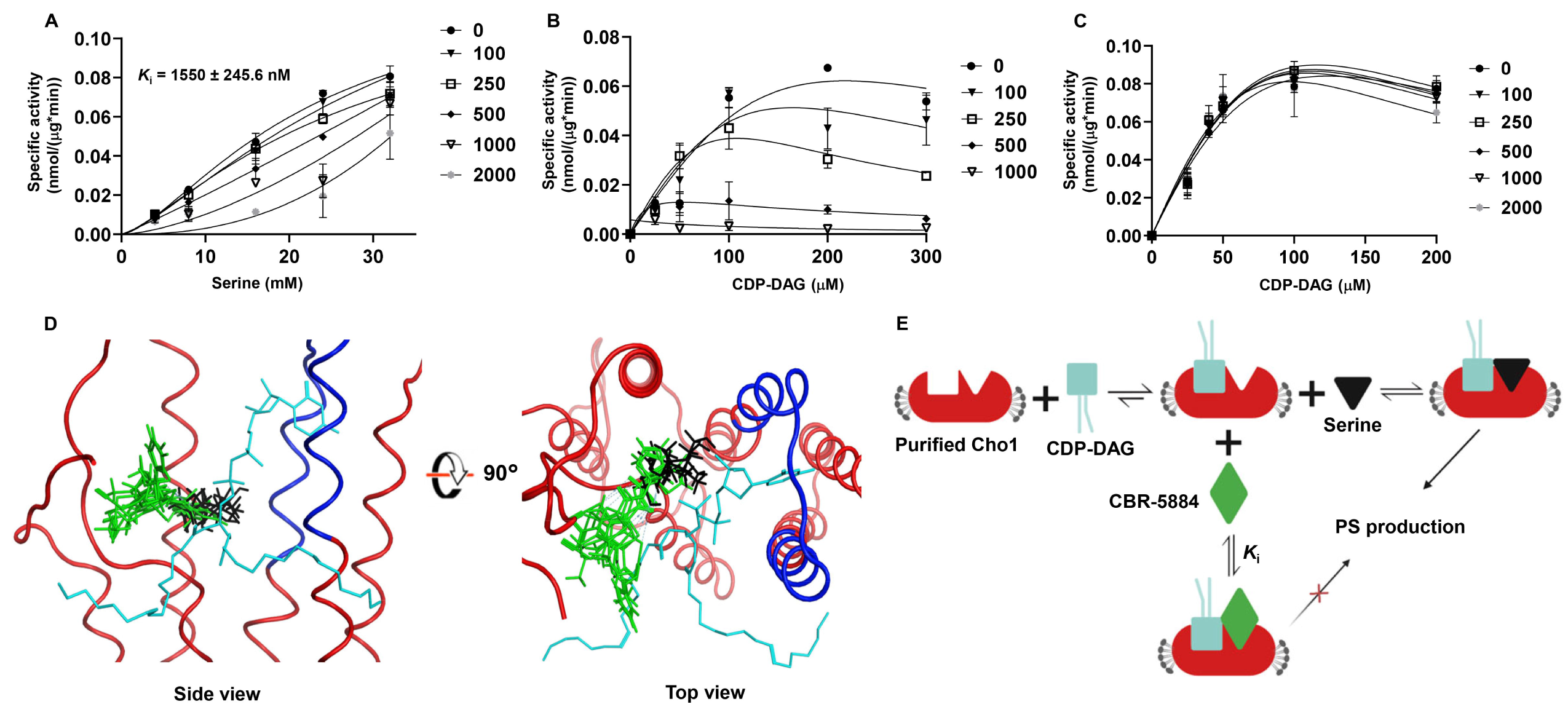
CBR-5884 may function as a competitive inhibitor occupying the serine binding site of Cho1. (A) Kinetics curve for serine in the presence of CBR-5884. CDP-DAG was kept constant at 200 μM (4.8 mol %), and the specific activities of purified hexameric Cho1 were plotted against various serine concentrations (4, 8, 16, 24, 32 mM) in the presence of 0, 100, 250, 500, 1000, 2000 nM CBR-5884. The curves best fit for competitive inhibition with a *K*_i_ value of 1550 ± 245.6 nM. The dots in all curves represent the mean values of two replicates, and the error bars are ± standard deviation (S.D.) values. **(B)** Kinetics curve for CDP-DAG in the presence of CBR-5884. Serine was kept constant at 20 mM, and the specific activities of purified hexameric Cho1 were plotted against various CDP-DAG concentrations (25, 50, 100, 200, 300 μM, 4.2 – 5.0 mol %) in the presence of 0, 100, 250, 500 and 1000 nM CBR-5884. **(C)** Kinetics curve for CDP-DAG in the presence of CBR-5884 with 32 mM serine. Specific activities of purified hexameric Cho1 were plotted against various CDP-DAG concentrations in the presence of 0, 100, 250, 500, 1000 and 2000 nM CBR-5884. **(D)** Computational docking of serine and CBR-5884 into the Cho1 AlphaFold structure. The top 5 poses of CBR- 5884 (green) and serine (black) are shown within the active site of Cho1. CDP-DAG is shown as cyan and the conserved CAPT motif of Cho1 is highlighted in dark blue. **(E)** A model for the inhibition mechanism of CBR-5884 to Cho1. Cho1 follows a sequential bi- bi reaction, in which it has to bind CDP-DAG prior to serine for catalysis. In the presence of CBR-5884, CDP-DAG-bound Cho1 could either bind serine for a reaction or CBR- 5884 for no reaction.

To gather more insight for the serine competition, we computationally docked CBR-5884 into the active site of *C. albicans* Cho1. Since fungal PS synthases follow the ordered sequential bi-bi reaction mechanism where Cho1 binds CDP-DAG before serine for catalysis (25), we first generated a predicted CDP-DAG-bound *C. albicans* Cho1 structure by superposing the *C. albicans* Cho1 AlphaFold model on the CDP-DAG-bound PS synthase from *Methanocaldococcus jannaschii* (PDB: 7B1L) (Figure S4A) (32). The CDP-alcohol phosphotransferase (CAPT) binding motif, which is known to bind CDP-DAG, is conserved and aligned between the *C. albicans* Cho1 and *M. jannaschii* PS synthase structures, so we hypothesized that the CDP-DAG from *M. jannaschii* PS synthase interacts with *C. albicans* Cho1 in a very similarly manner, and thus is incorporated in the *C. albicans* Cho1 model for docking. Next, we simulated and combined all the possible active site pockets from the CDP-DAG-bound *C. albicans* Cho1, and docked CBR-5884 and L-serine into these possible sites (Figure S4B). A total of 20,000 initial poses of CBR-5884 and serine were generated and then refined to 5 top poses with the highest docking scores (Figure 6D). All five CBR-5884 poses have overlap with the serine poses, with a higher docking score of -11.48 ± 0.12 kcal/mol compared to the serine docking score of -6.72 ± 0.08 kcal/mol. This, corroborates the low *K*_i_ value and suggests that CBR-5884 likely competes with serine to occupy the serine binding pocket in Cho1. Here, we generated a scheme for CBR-5884 inhibition (Figure 6E). Following the first step in the sequential bi-bi reaction where Cho1 binds CDP-DAG, either CBR-5884 or serine can dock into the active site. The catalysis will occur if serine enters, and will not if CBR-5884 occupies the site. Since CBR-5884 is favored by Cho1, serine can only outcompete CBR-5884 at a high concentration.

## Discussion

Here, we adapted a malachite green-based nucleotidase-coupled assay to screen for and identify inhibitors targeting *C. albicans* Cho1 (49). Since the amount of phosphate released is directly proportional to the CMP, and thus PS produced in the reaction, this method can be used to measure Cho1 activity in real time (Figure 1C). It is worth mentioning that besides CMP, the nucleotidase CD73 is known to cleave AMP to release phosphate in various studies (60–62), and given that this assay is suitable for 384- well plates or even 1536-well plates, it may potentially be applied to any enzyme producing AMP/CMP.

Ebselen, LOC14 and CBR-5884 stood out among seven Cho1-specific inhibitors due to their inhibitory effects on the *C. albicans* cell growth (Figure 3). Ebselen is an organo-selenium compound originally developed as a glutathione peroxidase mimic that acts on the cholesterol ester hydroperoxides and phospholipid hydroperoxides (63), and it has garnered significant attention in recent years due to its diverse therapeutic applications due to its anti-inflammatory, antioxidant and anticancer activity (64–69). In addition, ebselen has exhibited notable *in vitro* and *in vivo* antifungal activity against a range of fungal pathogens, including *Candida* spp., *Fusarium* spp., *Aspergillus fumigatus* and *Cryptococcus neoformans* (70–72). Studies have shown that ebselen effectively inhibits fungal growth by targeting many key enzymes (73–78). Here, we have demonstrated again that ebselen inhibits wildtype *C. albicans* growth (Figures 3&4). The inhibition of purified Cho1 by ebselen is potent, with an IC_50_ of 11.1 μM (Figure 2C), but the inhibition on Cho1 is very likely not the main cause for ebselen’s inhibition of *C. albicans* growth, since ethanolamine cannot bypass the drug (Figure 4). Currently, we do not know the mechanism of ebselen’s inhibition on Cho1, but given the tendency for ebselen to interact with cysteine residues (79), we hypothesize that ebselen might interact with residue C182, located in the putative serine-binding site of Cho1 (26), thus disrupting activity. Ebselen’s promiscuous nature limits its clinical applicability.

Conversely, the antifungal effects of LOC14 and CBR-5884 are less well studied. LOC14 is a potent, non-covalent and reversible inhibitor for protein disulfide isomerase that has a neuroprotective effect in corticostriatal brain culture (80). In addition, LOC14 displayed promising effects against Huntington’s disease (81) and can be used in a new anti-influenza therapeutic strategy (82). Here, we showed that LOC14 inhibits Cho1 activity (Figure 3), however, the *in vivo* inhibition is not likely conveyed through Cho1 (Figure 4). In contrast, CBR-5884 is an inhibitor of phosphoglycerate dehydrogenase, blocking *de novo* serine synthesis in cells, and is selectively toxic to cancer cell lines with high serine biosynthetic activity against melanoma and breast cancer lines (54). Here, CBR-5884 was shown to not only inhibit purified Cho1 (Figures 2&6), but also inhibit live cell growth by acting on the Cho1 *in vivo* (Figures 3-5). To our knowledge, this is the first paper showing the antifungal effects of LOC14 and CBR-5884.

CBR-5884 was then determined to be a competitive inhibitor of Cho1 with a *K*_i_ of 1550 ± 245.6 nM via kinetic analysis (Figure 6). Interestingly, in non-saturating serine concentrations (Figure 6B), the kinetic curves decreased in height and shifted to the left in the presence of increasing concentration of CBR-5884, indicating a decreasing *K*_m_ and *V*_max_, mimicking an uncompetitive inhibition where the inhibitor only binds to substrate-bound enzyme complex and thus depletes its population. This suggests that CBR-5884 is able to compete with serine by binding CDP-DAG-bound Cho1, and it cannot compete with CDP-DAG to bind to empty Cho1. The *V*_max_ of Cho1 in this study is estimated to be 0.128 ± 0.029 nmol/(μg * min), which is close to 0.088 ± 0.007 nmol/(μg*min) as described in (33). Also, it is worth mentioning that the curves where CDP-DAG was held constant and serine was varied follow a sigmoidal shape, which is consistent with previous finding that serine-binding may be cooperative (33), but the underlying mechanism is not clear. One small discrepancy has to be pointed out that the *K*_half_ of serine in this study is determined to be 21.88 ± 6.97 mM, which is five time higher than the 4.17 ± 0.45 mM from (33). The increased *K*_half_ may be explained by the presence of DMSO in this study, as DMSO has been shown to increase *K*_m_ in some enzymes (83–85).

CBR-5884 and serine were also docked onto the predicted CDP-DAG-bound Cho1 structure, and CBR-5884 was found to overlap with serine in the pocket (Figure 6D). A detailed ligand interaction map has shown that CBR-5884 almost shielded the β- phosphorus of the CDP-DAG where the nucleophilic attack occurs (30, 35), and also some CBR-5884 poses directly interact with residues R186 and F190 in Cho1 (Figure S5), which are part of the putative serine-binding site and are shown to be essential for Cho1 activity (26). This indicates again that CBR-5884 inhibits Cho1 by competing with serine.

Our data strongly suggest that CBR-5884 inhibits Cho1 *in vivo*, however, Cho1 may not be the only cellular target of CBR-5884. This compound causes cells to become more unmasked than the *cho1*ΔΔ mutant (Figure 5B). This could be due to sudden drop in PS caused by the compound compared with the *cho1*ΔΔ mutant, which has adjusted to the change. However, it could also be due to inhibition of another target. In addition, in TLC analysis, it was revealed that CBR-5884 decreased PS and also impacted relative PE levels similarly to that seen in a *cho1*ΔΔ mutant (Figures 5E&5F). However, the CBR- 5884 compound caused a greater decrease in PE levels in the *cho1*ΔΔ mutant than that observed in wild-type or untreated *cho1*ΔΔ cells. The compound also decreases growth of the *cho1*ΔΔ strain in rich YPD media (Figure S6), suggesting that it has an additional target besides PS synthase. Since the *cho1*ΔΔ strain cannot use PS as the precursor to make PE, the PE is made primarily via the Kennedy pathway, which requires CDP-ethanolamine phosphotransferase (14, 86). Cho1 and CDP-ethanolamine phosphotransferase belong to the same protein family and both use the CDP-alcohol phosphotransferase (CAPT) binding motif (29, 87, 88), so it is possible that CBR-5884 also inhibits CDP-ethanolamine phosphotransferase activity. Interestingly, PI synthase also has the CAPT motif and binds CDP-DAG (89, 90), but the PI levels were not strongly affected (Figure 5E&F). Thus, the impact would be more specific to ethanolamine phosphotransferase in this case.

This cross-reactivity could potentially be addressed by medicinal chemistry to synthesize analogs more specific for Cho1 and act at lower IC_50_ so such a high concentration is not required, which could decrease off-target effects. Moreover, the compounds could potentially be made more specific for Cho1. Importantly, we have novel proof of principle for successful pharmacological inhibition of a uniquely fungal enzyme central to phospholipid metabolism. Future rational drug design study will optimize CBR-5884 to be more specific for Cho1, as well as increase the solubility and potency of the compound in live cells.

## Materials and Methods

### Strain construction and media

This study used a *C. albicans* strain derived from SC5314 that was disrupted for *CHO1*, but had the gene complemented back with an affinity-tagged version. This strain *cho1*ΔΔ *P_TEF1_-CHO1-ENLYFQG-HAx3-HISx8*, was made in this study and used to solubilize and purify Cho1. This strain expressed a Hisx8-tagged Cho1 protein from a strong constitutive promoter *P_TEF1_*. To create this strain, a plasmid was generated that carried the tagged *CHO1* gene. This plasmid had a hygromycin B resistance gene, *CaHygB*, as a marker to go into strains. To make the plasmid pKE333, first the *CaHygB* marker cassette was amplified from the *CaHygB*-flipper plasmid (91) using primers YZO113 & YZO114 having *NheI* and *MscI* cut site, respectively (Table 1). The *CaHygB* cassette was ligated into pKE4 plasmid (50) that had been digested with *NheI* and *MscI*, to create the plasmid pKE4 - *CaHygB* (pKE333). Then, the *CHO1-ENLYFQG-HAx3- HISx8* gene, with the 3’UTR region was amplified from pYZ79 (33) using primers YZO110 & YZO111 and ligated into pKE333 cut with *ClaI* and *MluI* to create the plasmid pYZ107. Plasmid pYZ107 was linearized with *PmlI* restriction enzyme (within the *P_TEF1_* sequence) and electroporated into the *cho1*ΔΔ strain (14). Transformants were selected on YPD plates containing 600 μg/ml hygromycin B. Colony PCR was performed on six candidates for each gene construct to ensure the successful integration under the *P_TEF1_* promotor on the chromosomal DNA, and no spurious mutations occurred during the transformation. Media used in this study include YPD (1% yeast extract, 2% peptone, 2% dextrose) and minimal medium (0.67% yeast nitrogen base W/O amino acids, 2% dextrose ± 1 mM ethanolamine).

**Table 1.**
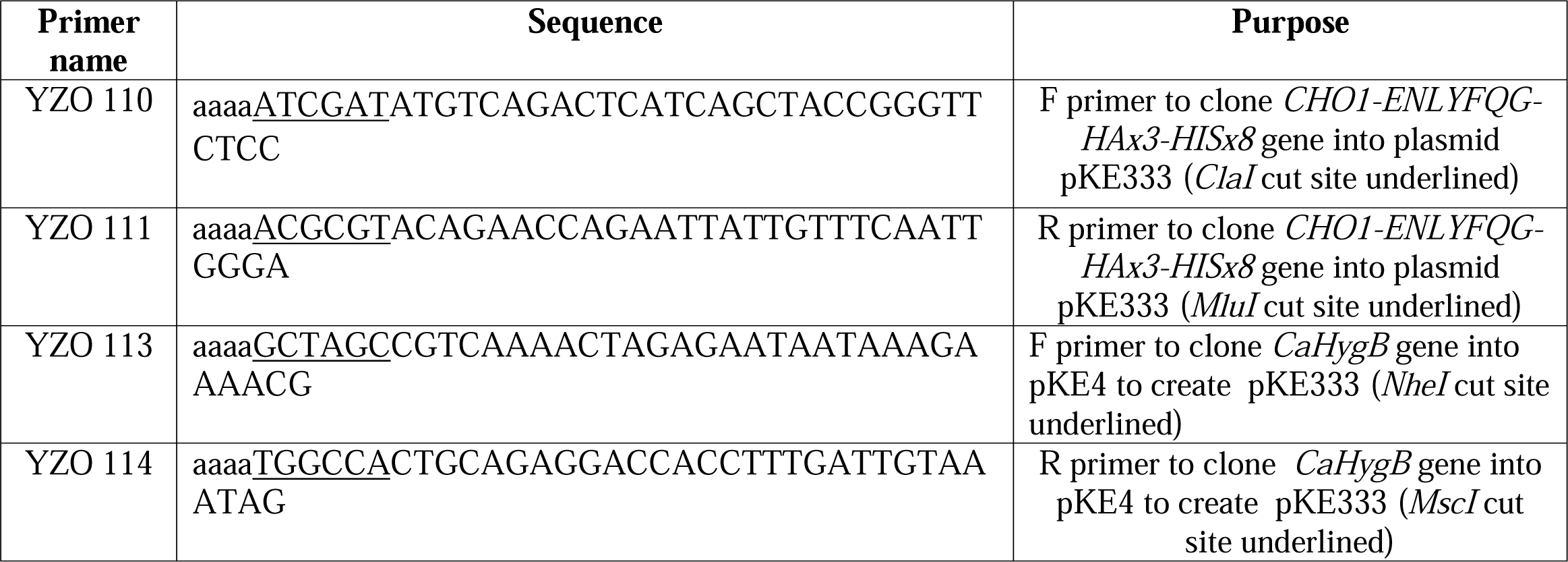
Primers used in this study.

### Cell lysis, protein solubilization and purification

The *C. albicans* strain with His-tagged Cho1 expressed from *P_TEF1_* was grown in YPD till the OD_600_ reached between 7 and 8, then cells were lysed using a French press, as described in (47). Crude membranes were collected and solubilized with 1.5% digitonin as described in (33). His-tagged Cho1 was first purified via gravity affinity chromatography as describe in (33), and was further cleared by size exclusion chromatography (Superdex 200 10/300 GL column (Cytiva) attached to an NGC chromatography system Quest 10 plus (Bio-Rad)). The column was equilibrated with 1.5 column volumes of H_2_O and 1.5 column volumes of elution buffer (50 mM Tris-HCl (pH=8.0) + 0.04% digitonin) with a flow rate of 0.5 mL/min. Samples loaded onto the above column consisted of a concentrated Cho1 eluted from affinity chromatography that was filtered through 0.22 μm filters that was then manually injected in the sample loop. The injected sample was eluted at a flow rate of 0.4 mL/min. Fractions containing Cho1 were pooled and subjected to AcTEV treatment, and then run through another round of affinity chromatography to remove impurities as described in (33). The resulting sample was loaded and checked on the blue-native PAGE for purity, oligomer state and homogeneity.

### High-throughput malachite green screen

The Bioactives library was purchased from Selleck in 2014, and the remaining libraries (anti-infectives, FDA-approved Drug, and mechanism of action collections) were assembled from compounds available at the Chemical Biology and Therapeutics Department at St. Jude Children’s Research Hospital (PMID: 33453364). All compounds were dissolved in DMSO and 100 nL was transferred to a 384-well clear bottom plate (ThermoFisher Scientific, cat# 265203) using a Beckman Echo 650 acoustic liquid handler. Equal volumes of either the selective CD73 nucleotidase inhibitor AB680 at a final concentration of 1 μM (MedChemExpress, cat# HY-125286) or DMSO were used as positive and negative controls, respectively. A total of 30-35 ng of purified Cho1 protein was used in each reaction in 50 mM Tris-HCl (pH=8.0) for primary screening, combined with 100 μM CDP-DAG (Avanti, cat# 870510), 5 mM serine, 0.4 ng CD73 nucleotidase (R&D systems, cat# EA002), 1 mM MnCl_2_, 0.1% APX-100 and 0.1% digitonin, in a total volume of 10 μL. Counter screen reactions were set up by replacing only the purified Cho1 with 150 μM CMP, with all other components remaining constant. The 10 μL primary and counter screening reactions were delivered to each well of 384- well assay plates containing pre-aliquoted compounds from the library using a Multidrop Combi (ThermoFisher Scientific). This resulted in a DMSO concentration of 1%. After media transfer, assay plates were incubated for 3 hours at 30 °C. Plates were then removed and 32 μL of malachite green mixture (6 μL malachite A (2% (w/v) ammonium molybdate and 20% (v/v) sulfuric acid in H_2_O) + 20 μL H_2_O + 6 μL malachite B (0.1% (w/v) malachite green oxalate and 0.5% (w/v) polyvinyl alcohol in H_2_O)) were added to each well via a MultiDrop Combi and incubated for an additional 10 minutes at room temperature. After incubation, plates were briefly centrifuged at 500 xg and absorbance at 620 nm was then measured using a Cytation7 plate reader (Biotek, Winooski, VT). Raw absorbance values of the compounds were normalized to DMSO (0% inhibition) and AB680 (100% inhibition) from both primary and counter screens, and % Δinhibition was calculated by subtracting % inhibition from the primary screen from the % inhibition from the counter screen. Z-factors for each plate were calculated using the in-house program RISE (Robust Investigation of Screening Experiments). All non-PAINs compounds that had >80% Δinhibition progressed to dose-response testing (82 total compounds). Dose-response experiments were performed in triplicate as described above using a 10-point, threefold serial dilution with the top concentration for each compound tested being 200 μM (Range: 0.010161-200 μM). The absorbance at 620 nm was then measured using a PHERAstar FS multilabel reader (BMG, Cary, NC). Raw values were once again normalized to DMSO (0% inhibition) and AB680 (100% inhibition) and Z-factors for each plate were calculated using RISE. The concentration of test compounds that inhibited Cho1 by 50% (IC_50_ value) as measured by the malachite green assay was computed using nonlinear regression-based fitting of inhibition curves using [inhibitor] vs. response-variable slope model in GraphPad Prism version 9.5.0 (GraphPad Software, La Jolla California USA).

### Assay for metabolic labeling of PS using probe C-L-SerN_3_

The C-L-SerN_3_ probe ((*S*)-1-((3-azidopropyl)amino)-3-hydroxy-1-oxopropan-2- aminium chloride) was synthesized as described in (57). Cells grown overnight in YPD were washed three times in H_2_O and inoculated into minimal media at a starting OD600 of 0.05. Cells were shaken for 5 hours before 1.5 mM C-L-SerN_3_ probe was added, along with 170 μM CBR-5884 or an equivalent volume of DMSO. Cells were then incubated for another 5 hours before being washed three times with H_2_O, and 5% BSA in 1xPBS was used to treat cells for 20 min. The cells were then washed three times with 1xPBS, and then resuspended to OD_600_ of 0.6 in 1 mL PBS + 1 μM AZDye 488 DBCO (Click chemistry tools, cat# 1278-1). Cells were covered with aluminum foil and rocked for 1 hour, and the dye was removed by pelleting the cells at 5,000 xg followed by washing with shaking at 1,000 rpm three times in 1xPBS, and resuspended in 200 μL Fluoromount-G mounting medium (cat# 00-4958-02). For each treatment, 10 μL of cell was added to a glass slide and 3 μL fresh Fluoromount-G mounting medium was used to mix the sample. Then, a Leica SP8 white light laser confocal microscope was used for imaging. The samples were excited using light at a wavelength of 488 nm, and the resulting fluorescence was captured within the range of 498 to 550 nm using a HyD detector. The settings for laser strength, gain, and offset were maintained consistently throughout the experiment. Images of treated cells were taken after applying a zoom factor of 3. A total of 40 cells from at least 10 images were used for the quantification in ImageJ software.

### Fluorescence imaging of unmasked **β**(1–3)-glucan

Wildtype and *cho1*ΔΔ cells were grown in YPD overnight (∼16 hours), and back diluted to fresh YPD with OD_600_ of ∼0.1. The cells were then shaken at 225 rpm for 3 hours before 170 μM CBR-5884 or equivalent DMSO were added. The cells were further shaken for 30, 60, 120 mins before the exposed β(1–3)-glucan staining. The cells were stained as previously described (55, 92) with the exception that goat anti-mouse antibody conjugated to Alexa Fluor® 488 (Jackson ImmunoResearch) was used as the secondary antibody. For imaging, *Candida* cells were resuspended in 200 μL of Fluoromount-G mounting medium and visualized with a Leica SP8 white light laser confocal microscope. The pictures were taken through Leica Application Suite X office software.

### MIC plates tests and growth curves

MIC plate assays and growth curves were conducted in minimal media or minimal media supplemented with 1 mM ethanolamine (ETA). Wildtype SC5314 and *cho1*ΔΔ strains were grown in YPD overnight and washed three times with H_2_O before inoculation. The starting OD_600_ is 0.05 for both plate assays and growth curves. For MIC plates, both wildtype and *cho1*ΔΔ strains were grown in flat-bottom 96-well plates at a final volume of 200 μL, in the presence of different compounds or DMSO as indicated at 30°C. A Leica light inverted microscope was then used to visualize growth after 24-hour incubations. In the meantime, cells grown in different compounds were subjected to live cell counting. Cell cultures from flat-bottom 96-well plates after 24-hour incubation were diluted 100 to 10,000 times before plating on the minimal media, and the total colony forming unit (CFU) was kept within 200 for each plate. A total of six replicates were done in each condition.

The growth curves with different treatments were measured from 2 to 30 hours with OD_600_ measurement every two hours. A final concentration of 170 μM CBR-5884, 430 μM LOC14 and 5 mM serine was added into each group as indicated. Each curve was plotted in duplicate with six biological replicates. Growth rate (h^-1^) and lag time (h) were calculated using GraphPad Prism software.

### PS synthase assay

Enzymatic activity of Cho1 in the presence of the compounds was measured using the radioactive PS synthase assay. For purified Cho1, the procedure was fully described in (33). Briefly, 1-2 μg of purified Cho1 was added in the reaction containing 50 mM Tris-HCl (pH = 8.0), 1 mM MnCl_2_, 0.1% Triton X-100, 0.04% digitonin, 0.1 mM (4.7 mol %) 16:0 CDP-DAG (Avanti, cat# 870510) and 0.5 mM L-serine (spiked with 5% (by volume) L-[^3^H]-serine (15 Ci/mmol)) at a total volume of 100 μL, in the presence of 100 μM compounds or equivalent DMSO solvent. The reaction was conducted at 30 °C for 30 min and synthesized [^3^H]PS was measured using a liquid scintillation counter. For crude membrane samples, crude membrane preps from wildtype and *cho1*ΔΔ strains were collected and assayed as described in (33) with the exception that 2 mM L-serine (spiked with 5% (by volume) L-[^3^H]-serine (15 Ci/mmol)) was used in each reaction. A final concentration of 1 mM compounds was used for the crude membrane samples.

For the kinetic curves, the specific activity was measured in the reaction containing 50 mM Tris-HCl (pH = 8.0), 1 mM MnCl_2_, 0.025-0.3% Triton X-100 and 0.033-0.07% digitonin with 0.75 to 1.5 μg purified Cho1 protein. The concentrations of 18:1 CDP-DAG (Avanti, cat# 870520) and serine were indicated in the graphs, and the mol % of CDP-DAG was kept between 4.2 – 5.0% for the curve where CDP-DAG was varied (Figures 6B & 6C) and at 4.8% (200 μM) for the curve where serine was varied (Figures 6A). The concentrations of CBR-5884 used in each reaction were indicated in the graph, and CBR-5884 was incubated with purified Cho1 protein on ice for at least 2 hours before the addition of L-[^3^H]-serine. The reaction was stopped at 20, 40 and 60 min, and specific activity was calculated based on the slope of linear PS production, representing the initial velocity.

### Thin layer chromatography

Wildtype SC5314 and *cho1*ΔΔ strains grown overnight in YPD were inoculated into fresh YPD at OD_600_ of 0.1 and were shaken for another 3 hours. Then, 170 μM CBR- 5884 or equivalent DMSO solvent was added to both wildtype and *cho1*ΔΔ cultures, and cells were shaken for another 2 hours. Then, cells were washed with 1xPBS three times and normalized to a total OD_600_ of 1. The phospholipids were extracted with hot ethanol method as described in (14). A Whatman 250 μm silica gel aluminum backed plate was treated, and separation of phospholipids was carried out as described in (93).

Phospholipid standards PI, PE, PS and PC were purchased from Avanti. The quantification of the phospholipids was done in ImageJ software.

### Computational docking

The computational docking was conducted in Molecular Operating Environment software (MOE, Chemical Computing Group, Ltd, Montreal, Canada). A CDP-DAG-bound *C. albicans* Cho1 structure was generated by superposing the *C. albicans* Cho1 AlphaFold model on the CDP-DAG-bound PS synthase from *Methanocaldococcus jannaschii* (PDB: 7B1L) (32). Structures from CBR-5884 and serine were introduced into the structure, and the system was quickly prepped and energy-minimized for docking.

Potential docking sites were predicted by “site finder” function in MOE and all the sites having above 0 possibilities are combined for docking (Figure S4B). Both serine and CBR-5884 molecules were docked into the potential sites 20,000 times, and triangle matcher placement (scored by London dG) and rigid receptor refinement (scored by GBVI/WSA dG) were used to pick the top fives poses. The ligand interaction map was also generated in MOE.

### Statistical analysis and molecular weight (MW) estimation on the gels

All the statistical analyses were performed with GraphPad Prism 9.1 software.

The PS synthase activities were compared using ordinary (equal SDs) or Brown-Forsythe and Welch ANOVA tests (unequal SDs). Blue native PAGE and Coomassie Blue R-250 staining were conducted as described in (33) and all MW estimates were conducted in the band analysis tool of the Quantity One software (Bio-Rad).

### Data availability

The original contributions presented in the study are included in the article/Supplementary Material. Further inquiries can be directed to the corresponding author.

## Supporting information

Supplementary data combined

## Acknowledgments

The authors would like to thank Dr. Glen E. Palmer and Dr. Brian M. Peters from the University of Tennessee Health Science Center for their pKE4 and *CaHygB*-flipper plasmids. Reese Saho, a summer undergraduate student from Ohio Northern University, also helped with setting up the PS synthase assay. We also thank him for his work.

## Author contributions

Conceived and designed the experiments: YZ, GAP, JMR, REL, TBR. Performed the experiments: YZ, GAP, MMM, JM, JMR, EKP, JL, CFA. Analyzed the data: YZ, GAP, TBR. Contributed reagents/materials/analysis tools: JMR, REL, MDB, TBR. Wrote the original draft: YZ. Writing & Editing: YZ, GAP, PMO, CFA, TBR. All authors contributed to the article and approved the submitted version.

