## Supplementary data combined for "The small molecule CBR-5884 inhibits the *Candida albicans* phosphatidylserine synthase"

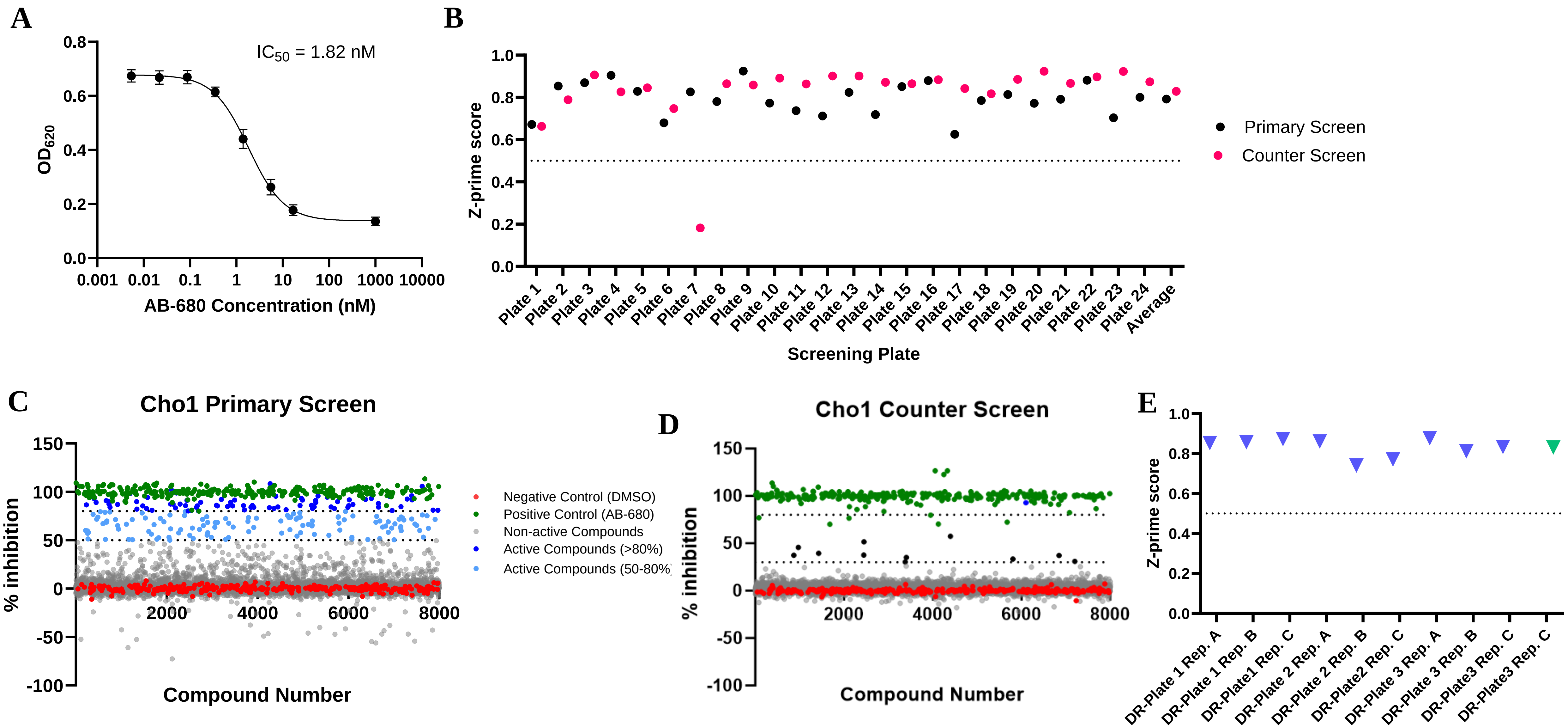


Figure S1. (A) Dose-response curve of AB-680 compound. The dots represent the mean of nine replicates, and the error bars are ± standard deviation (S.D.) values. (B) Z-prime scores of plates in the primary and counter screens. (C,D) The dot plots of % inhibition for all the compounds, including controls, used in the primary screen (C) and counter screen (D). Reactions with DMSO and AB-680 were used as 0% inhibition (negative) and 100% inhibition (positive) controls, respectively. The two dotted lines from the Y-axis indicate 80 and 50% inhibition, respectively. (E) Z-prime scores of plates in the dose-response curves for the second-round screen.

Figure S2. (A) Avasimibe, tideglusib and CBR-5884 precipitated at high concentrations; (B, C) The growth rate ($h^{-1}$) (B) and lag time (h) (C) were calculated from Figure 4D. Statistics were conducted using one-way ANOVA and Tukey's multiple comparisons test (ns=not significant, p > 0.05; *, 0.05> p > 0.01, ****, 0.0001> p).

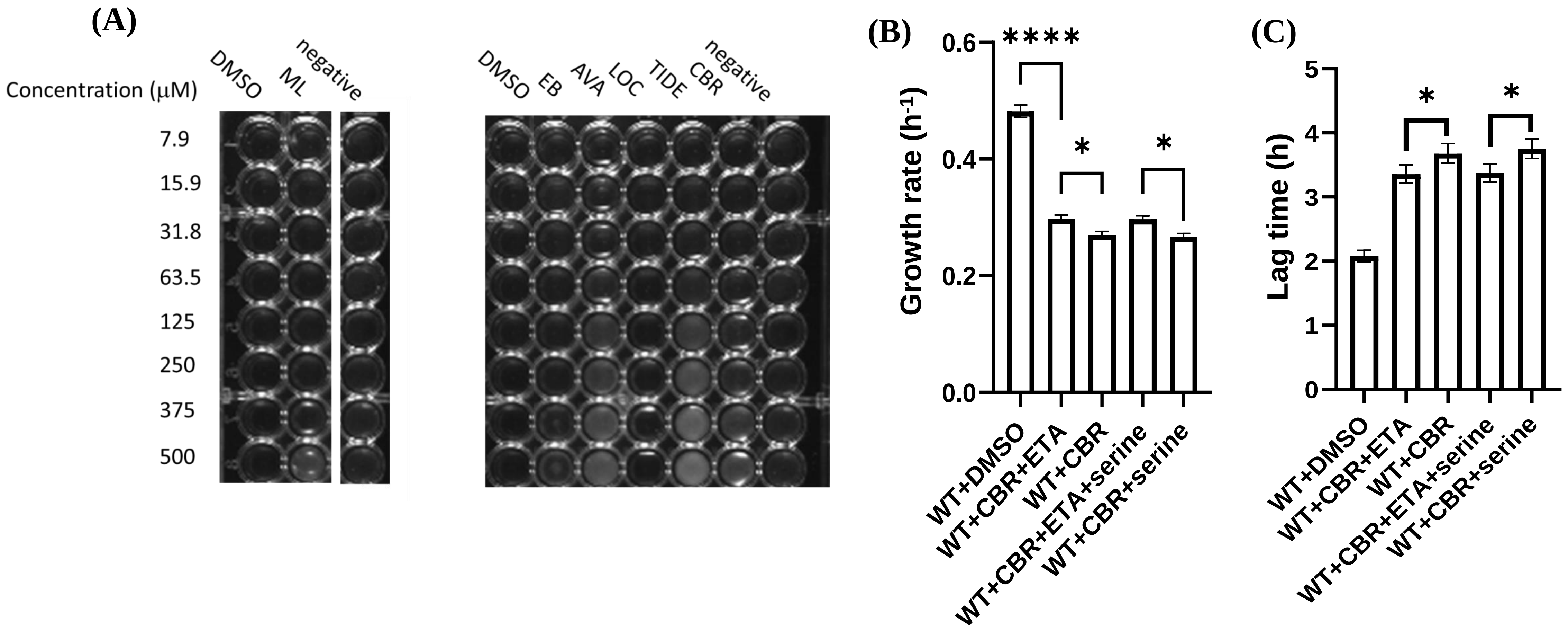

Figure S3. The PS synthase activity of crude membrane containing Cho1 was measured by L-[$^3$H]-serine incorporation into PS in the presence of different inhibitors at 1 mM, and are presented as nmol/(μg protein*min). Statistics were conducted using one-way ANOVA and Dunnett's T3 multiple comparisons test (**, $0.01 > p > 0.001$; ***, $0.001 > p > 0.0001$; ****, $0.0001 > p$). The activities were measured in duplicate with a total of six biological replicates as indicated. The bars represent the mean values and the error bars are ± S.D. values.

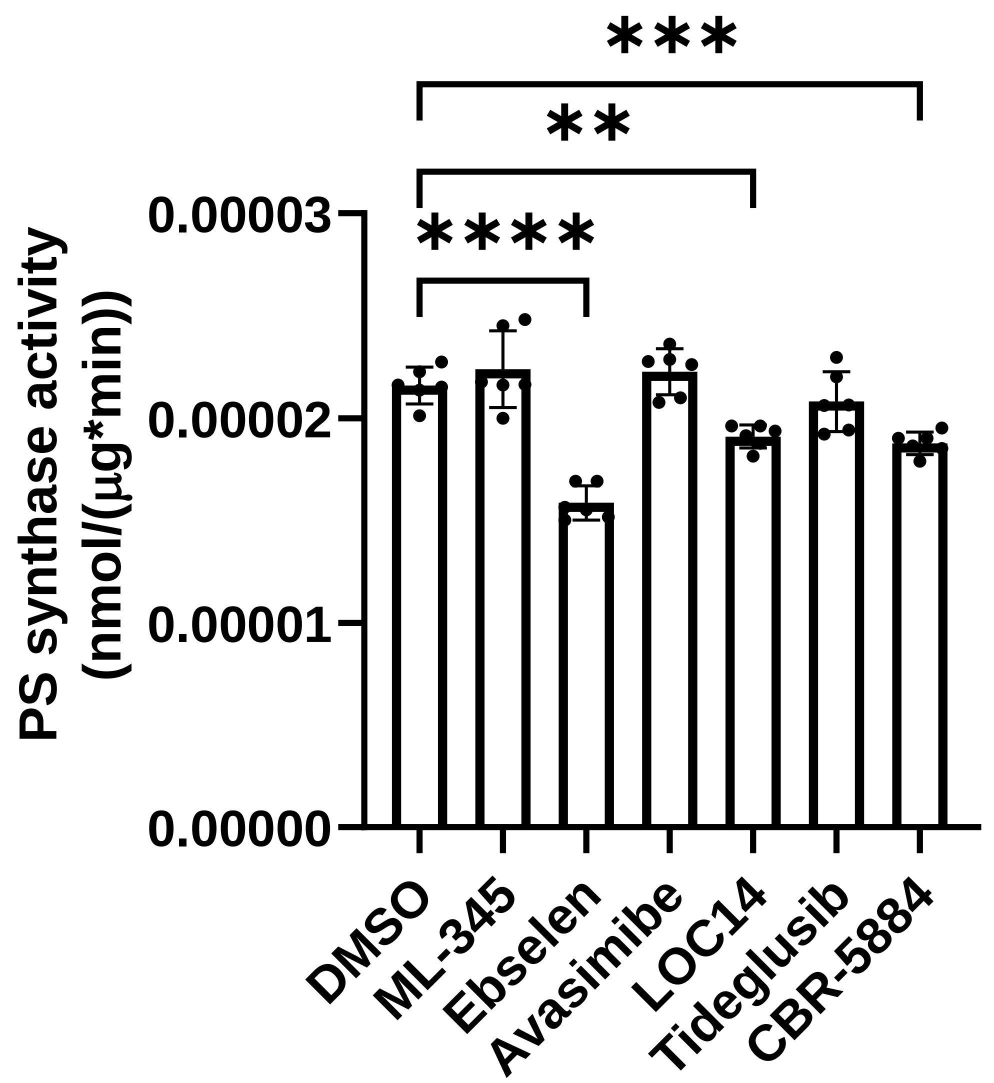

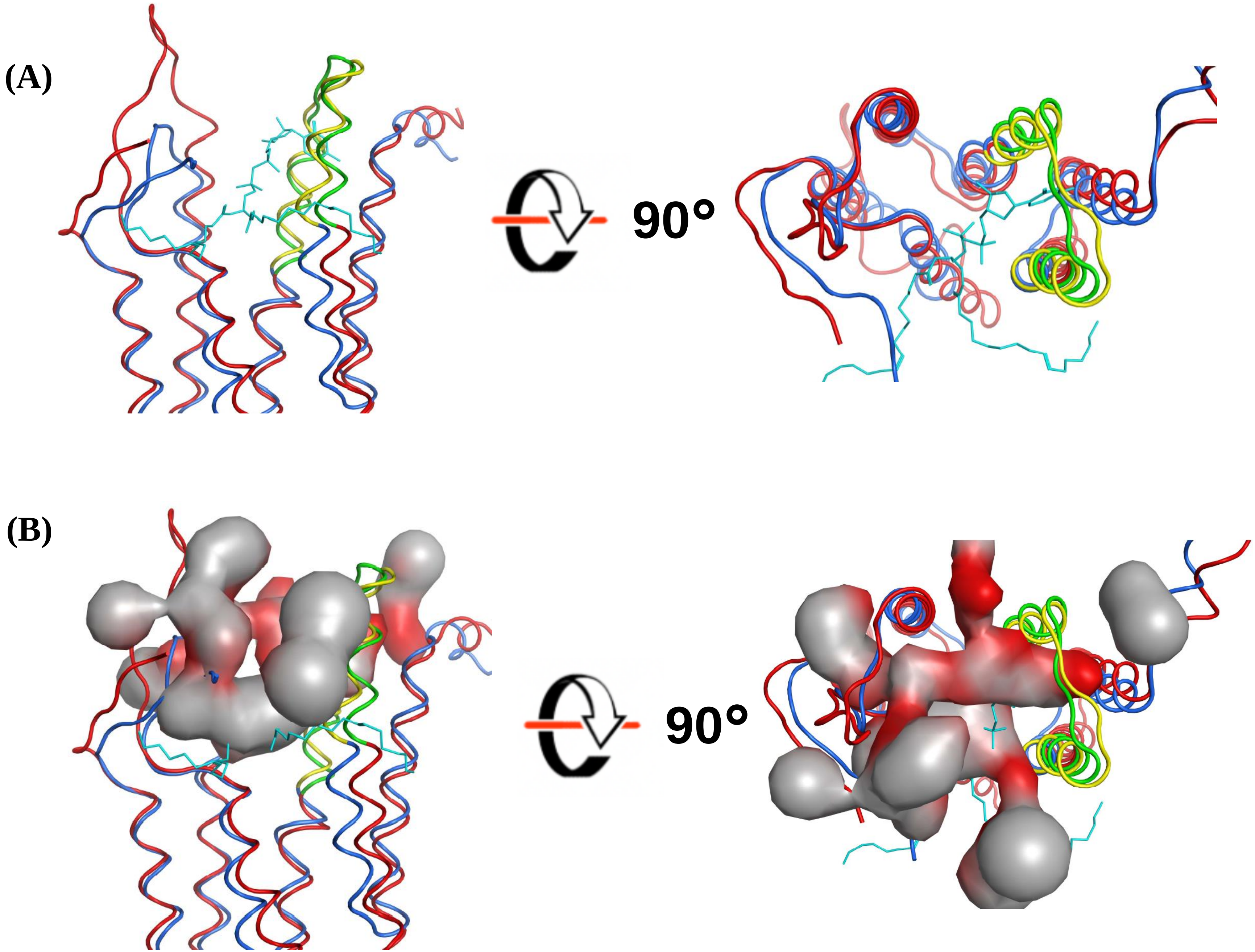


Figure S4. (A) Superposition of *C. albicans* AlphaFold Cho1 model (red) with *Methanocaldococcus jannaschii* PS synthase (blue) (PDB: 7B1L).The conserved CAPT site are highlighted as green and yellow, respectively. CDP-DAG (cyan) is *Methanocaldococcus jannaschii* PS synthase (PDB: 7B1L). (B) All the possible active site pockets predicted from *C. albicans* Cho1 using MOE are shown as pocket surfaces.

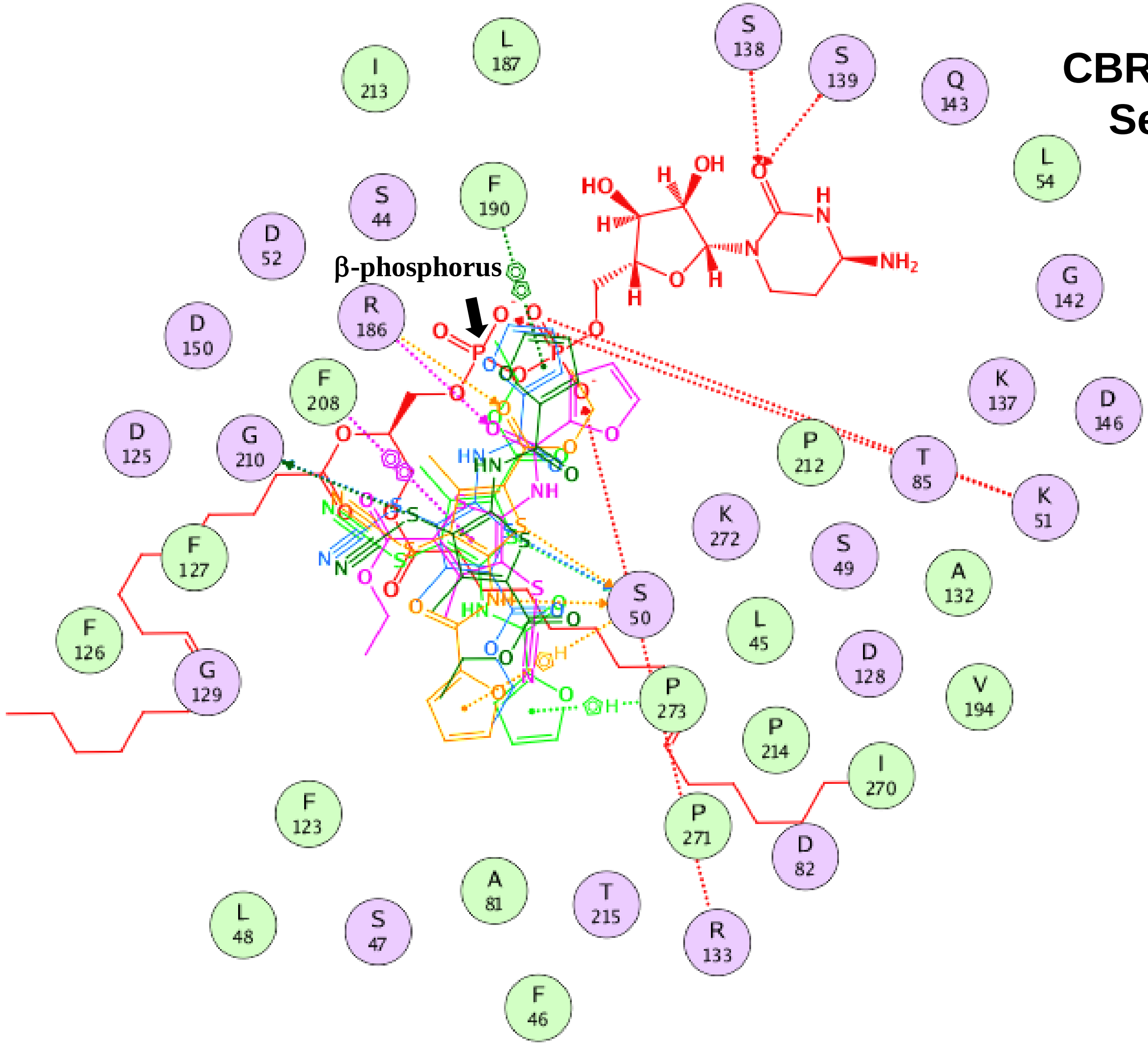


**CBR-5884 MOE(S) = -11.48 ± 0.12 kcal/mol**
**Serine MOE(S) = -6.72 ± 0.08 kcal/mol**

Figure S5. Overlay of ligand interactions between top five poses of CBR-5884 with *C. albicans* AlphaFold Cho1 model. The bound CDP-DAG is also shown and the β-phosphorus is indicated. Dotted lines indicate interactions. Docking scores MOE(S) of serine and CBR-5884 were shown.

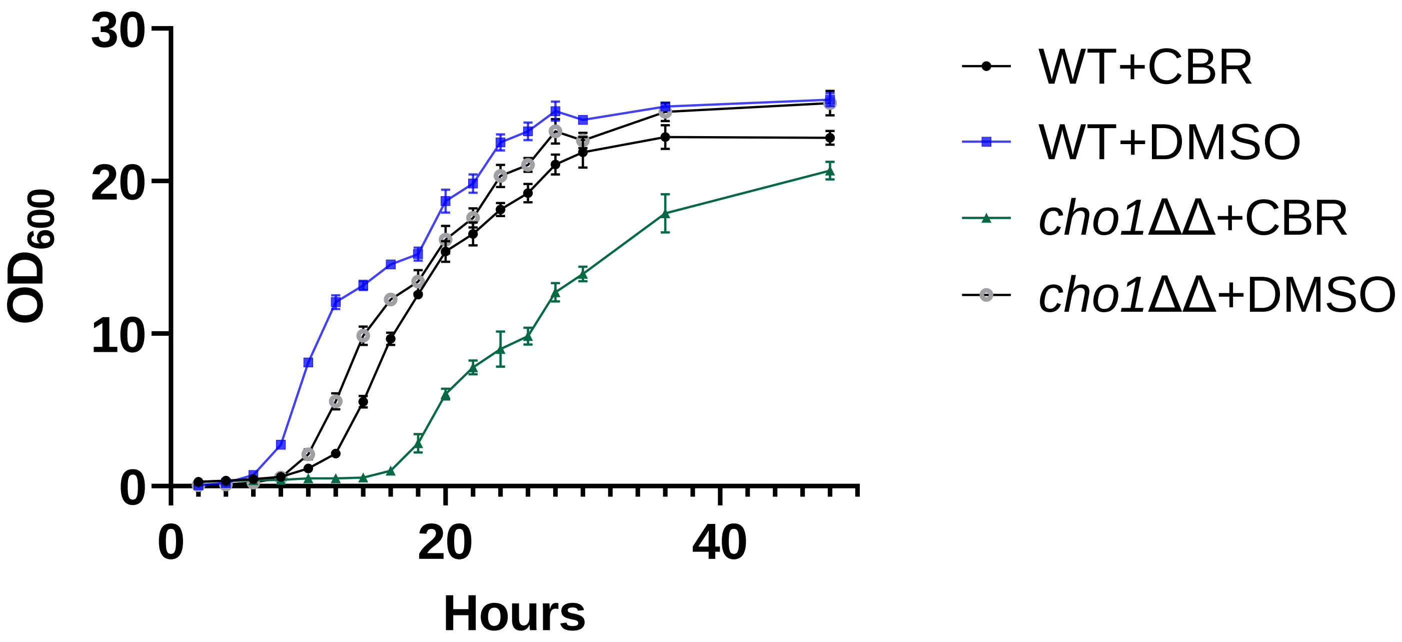


Figure S6. Growth curve of wildtype and *cho1*ΔΔ *C. albicans* strains in the presence of 170 μM CBR-5884 or equivalent DMSO in YPD from 0 to 48 hours. The dots represent the mean values of six replicates, and the error bars are ± standard deviation (S.D.) values
